# Prosaposin is cleaved into saposins by multiple cathepsins in a progranulin-regulated fashion

**DOI:** 10.1101/2024.02.15.580326

**Authors:** Molly Hodul, Courtney Lane-Donovan, Edwina Mambou, Zoe Yang, Aimee W. Kao

**Affiliations:** Department of Neurology, University of California, San Francisco, California, 94143, U.S.A.

**Keywords:** Prosaposin, progranulin, cathepsins, proteolytic enzyme, lysosome, sphingolipid, lipid metabolism

## Abstract

Prosaposin (PSAP) is a lysosomal protein that plays a key role in sphingolipid metabolism. PSAP is cleaved into four bioactive disulfide-rich peptides, saposins A, B, C, and D, that catalyze sphingolipidases to promote sphingolipid breakdown. Considering the key role of PSAP and saposins in sphingolipid metabolism and the existence of genetic mutations in PSAP associated with juve-nile-onset lysosomal storage and adult-onset neurodegenerative diseases, maintaining optimal levels of PSAP and saposins is crucial for proper lysosomal function and sphingolipid homeostasis. Despite this, the mechanism by which saposins are released from PSAP, and thus available to modulate sphingolipidases, sphingolipid homeostasis, and downstream lysosomal function, is not well understood. Here, we performed a comprehensive study to identify lysosomal enzymes which regulate prosaposin cleavage into saposins. *In vitro* cleavage assays identified multiple enzymes that can process human prosaposin into multi- and single-saposin fragments in a pH-dependent manner. We confirmed the role of cathepsins D and B in PSAP processing and showed that these and several previously unidentified lysosomal proteases (cathepsins E, K, L, S, V, G and AEP/LGMN) are able to process PSAP in distinctive, pH-dependent manners. In addition, we have demonstrated a novel role for progranulin (PGRN) in the regulation of PSAP cleavage. We found that PGRN and multi-granulin fragments (MGFs) directly regulate the cleavage of PSAP by cathepsin D. With this study, we have identified that multiple cathepsins, PGRN and MGFs work in concert to produce saposins under different conditions, which could present novel opportunities to modulate saposin levels in disease.

## INTRODUCTION

A key factor in neurodegenerative disease is the progressive deterioration of protein homeostasis, leading to macromolecule accumulation and cellular dysfunction. Given their role in the degradation, recycling and homeostasis of macromolecules, lysosomes have emerged as an important contributor to proteostasis. Moreover, lysosomal dysfunction is linked to juvenile-onset lysosomal storage and adult-onset neurodegenerative diseases [1]. Interestingly, perturbed lysosomal metabolism of a specific class of lipids, sphingolipids, is a critical risk factor for the onset and progression of neurodegenerative diseases [2-7].

Sphingolipid metabolism in the lysosome is driven by lysosomal enzymes called sphingolipidases. Sphingolipidase activator proteins, also known as SAPs or saposins, are bioactive disulfide-rich peptides that catalyze sphingolipidases to promote sphingolipid breakdown. Saposins regulate this process both through direct activation of sphingolipidases and by disrupting the intralysosomal membrane in a way that presents substrates to these enzymes [8-11]. Prosaposin (PSAP) is the evolutionarily conserved precursor for the four major sphingolipidase activator proteins, saposins A, B, C and D [12, 13].

As the precursor to saposins A-D, full-length PSAP is central to the sphingolipid catabolism pathway [12, 14-25]. Indeed, mice lacking PSAP show considerable accumulation of several sphingolipids [12, 26, 27]. Loss of PSAP in humans leads to Combined Saposin Deficiency, a fatal infantile lysosomal storage disorder with severe neurological pathology [28]. Interestingly, loss of each individual saposin leads to a different lysosomal storage disorder. Deficiency in SapA, SapB, or SapC leads to Krabbe disease [17, 29-31], metachromatic leukodystrophy [12, 32-36], or Gaucher disease [12, 24, 37, 38], respectively. In addition to lysosomal storage disorders, mutations in *Psap* are also linked to age-related neurodegenerative diseases such as Alzheimer’s Disease (AD) [39] and Parkinson’s Disease (PD) [40].

Despite the important roles for saposins A-D in sphingolipid metabolism and disease, the mechanism by which saposins are released from PSAP is poorly understood. As a lysosome resident protein, PSAP proteolysis likely occurs in this organelle [41, 42]. In fact, lysosomal proteases cathepsin D (CTSD) [43] and cathepsin B (CTSB) [44], have been shown to cleave PSAP into saposins. However, a comprehensive study of the PSAP cleavage capacity of lysosomal proteases has not been performed.

In addition to sphingolipidases, PSAP directly interacts and trafficks with another lysosomal pro-protein, progranulin (PGRN) [45, 46]. *Pgrn* loss of function mutations also lead to neurodegenerative diseases such as Frontotemporal Dementia (FTD) [47, 48]. Recently, PGRN has been implicated in disease-related dysfunction in sphingolipid homeostasis [49, 50]. In the lysosome, PGRN is cleaved by a subset of lysosomal proteases sequentially into multi-granulin fragment (MGF) intermediates and ultimately bioactive granulins [51]. There, PGRN and the MGFs promote the activation of CTSD [42, 52-55]. As CTSD has been shown to catalyze the cleavage of PSAP into saposins [43], this is one potential mechanism by which PGRN can indirectly influence lipid metabolism.

In this study, we set out to understand the processing and regulation of PSAP into saposins by lysosomal proteases. Using *in vitro* protease cleavage assays, we identified several lysosomal proteases that can process PSAP into trisaposins, disaposins and saposins and show that this processing occurs in a protease-specific and pH-dependent manner. Additionally, we found that progranulin and the MGFs have differing effects on PSAP cleavage, and these effects are mediated by CTSD.

## RESULTS

### Prosaposin is cleaved by a subset of lysosomal proteases *in vitro*

The endo-lysosomal compartment contains numerous proteolytic enzymes that are classified by the amino acid(s) in their active site, and are thereby assigned to aspartyl (*e*.*g*. CTSD), cysteine (*e*.*g*. CTSB), or serine (*e*.*g*. CTSA) families. *In vitro*, PSAP is cleaved into saposins by at least two cathepsins, CTSD [43] and CTSB [44]. Considering that individual proteases recognize distinct but overlapping amino acid motifs [56, 57], we postulated that several proteases are capable of processing of PSAP. To identify the lysosomal proteases that cleave PSAP, we performed an *in vitro* PSAP cleavage campaign with commercially available recombinant human lysosomal enzymes. Recombinant PSAP was incubated with individual lysosomal proteases at for one hour at 37°C, then cleavage products visualized via silver stain. As lysosomal proteases have distinct pH preferences that can be substrate-dependent [57-59], we performed the study across a range of pH settings (pH 3.5, 4.5, 5.5, and 7.4).

We first assayed the aspartyl proteases, cathepsins D and E. Like other proteases, CTSD is first translated as an inactive proenzyme (proCTSD) which is subsequently cleaved into active mature CTSD. As expected, we found that the inactive proCTSD was unable to cleave PSAP. In contrast, the mature active CTSD protealyzed PSAP into discrete peptides (Fig. 1A). However, despite being one of the only known cathepsins for PSAP [43], mature CTSD did not induce strong cleavage, even at its optimal pH (3.5). On the other hand, we found that under the same acidic conditions CTSE almost fully reduces PSAP to saposins (Fig 1A). Cathepsins D and E are highly related and recognize similar amino acid motifs [60]. Nonetheless, they exhibit different abilities to process PSAP into saposins, and there are subtle differences between preferred cleavage sites, which may explain the pronounced differences in activity we see here.

**Figure 1.**
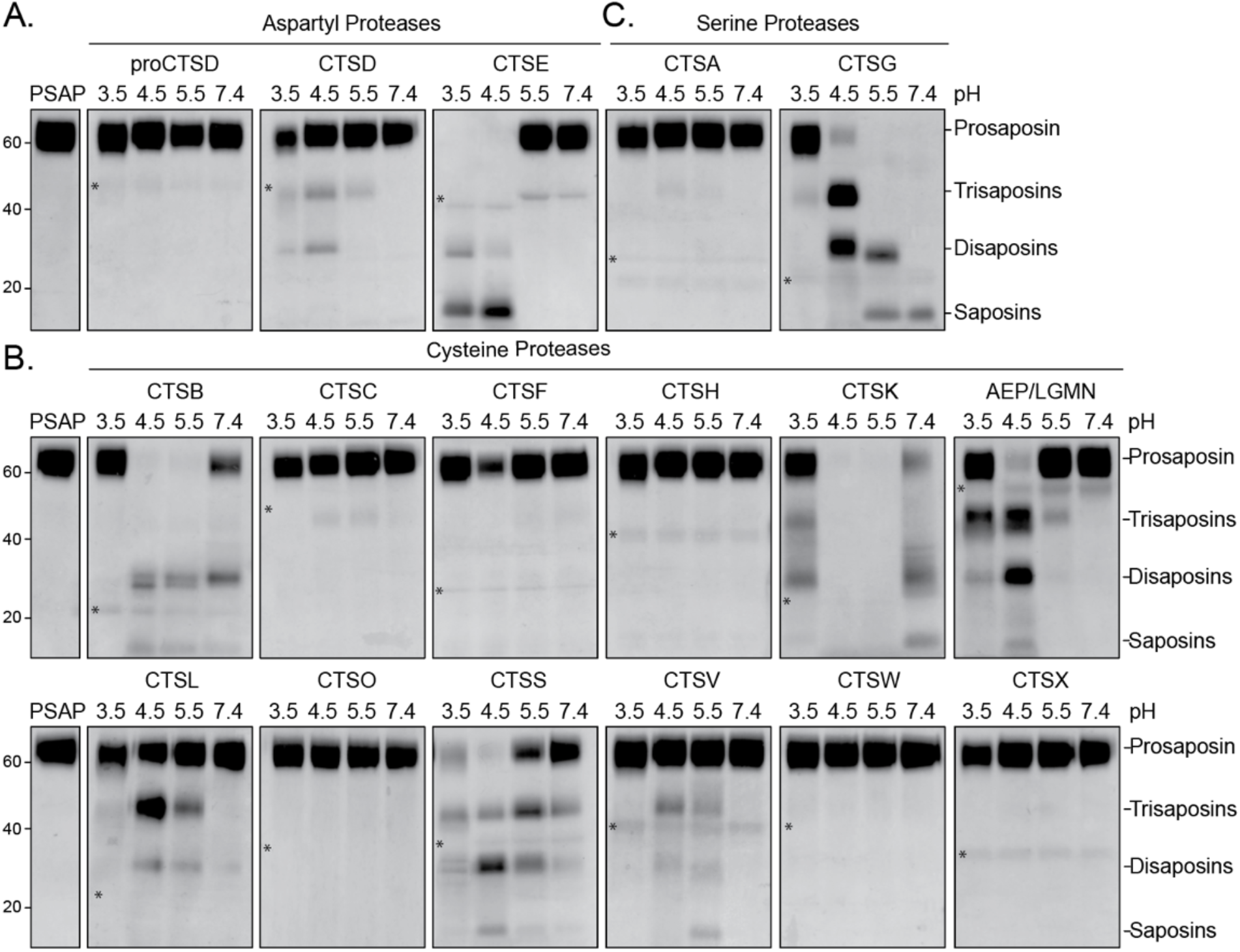
Prosaposin is cleaved into saposins by numerous cathepsins. Human recombinant prosaposin (PSAP, 400nM) was incubated at 37°C for 1 hour with various human recombinant cathepsins (400nM) under four pH conditions (3.5, 4,5, 5.5, and 7.4) and cleavage was determined via silver stain. *A*, Aspartyl proteases CTSD and CTSE, *B*, cysteine proteases CTSB, CTSC, CTSF, CTSH, CTSK, AEP/LGMN, CTSL, CTSO, CTSS, CTSV, CTSW, and CTSX, and *C*, serine proteases CTSA and CTSG were tested. * indicates the cathepsin band for respective experiments.

We next tested the cysteine proteases. Consistent with a previous study [44], we found that cathepsin B cleaves PSAP at a lysosomal pH of 4.5-5.5 (Fig 1B). Interestingly, this cleavage appeared to preferentially occur between SapB and SapC, as mostly disaposins were produced. In addition to CTSB, we found that cathepsins K, L, S, V, and AEP/LGMN also processed PSAP (Fig 1B). Each cathepsin generated different PSAP cleavage profiles, with products the size of trisaposins, disaposins, and/or saposins. CTSK was the most efficient cysteine protease, as it completely digested PSAP at pH 4.5-5.5 and produced cleavage products at pH 3.5 and pH 7.4. CTSS and AEP/LGMN also cleaved PSAP into several discrete fragments, almost completely clearing full-length PSAP at pH 4.5. Cathepsins L, V and F also proteolyzed PSAP, although not as efficiently as the other cysteine proteases. In contrast, cathepsins C, H, O, W and X did not digest PSAP at any pH tested (Fig. 1B).

Lastly, we tested the serine proteases cathepsins A and G. CTSG showed robust processing at pH 4.5, 5.5 and 7.4, while CTSA showed no cleavage of PSAP at any pH (Fig. 1C). Cleavage at a neutral pH is consistent with an extracellular role for CTSG [61]. Like CTSG, the cysteine cathepsins B, K, and S were also capable of processing PSAP at a neutral pH (Fig. 1B), suggesting that these proteases may process PSAP in both intracellular and extracellular compartments.

In summary, two aspartyl proteases (cathepsins D and E), six cysteine proteases (cathepsins B, K, L, S, V and AEP/LGMN), and one serine protease (cathepsin G) digested full-length PSAP *in vitro* in a pH-dependent manner. Since protease expression can be cell-type specific [51], these results suggest that PSAP processing may be carried out in a variety of ways, depending on the cell type and intravs. extracellular context.

### The efficiency of saposin production differed between cathepsins *in vitro*

The initial test for ability to cleave PSAP was performed over 1 hour. However, certain proteases (*e*.*g*. CTSE) completely processed PSAP while others (*e*.*g*. CTSD) did so incompletely. To better resolve the stepwise proteolysis of PSAP into tri-, di- or mono-saposins, the incubation time for each cathepsin was individualized. Thus, certain cathepsins were tested over a shorter incubation period (cathepsins E, B, K, S, and G) and others for a longer incubation (cathepsins D, L, V, and AEP/LGMN). Saposin fragment abundance was measured for those tested on the shorter incubation after 5 minutes, 15 minutes, and 30-minute incubations, and for those tested on the the longer incubation after 1 hour, 3 hour, and 6 hour incubations. Individual saposins are ∼9kDa, and could be detected with saposin-specific antibodies (Fig. 2A). At shorter time points, a second SapD band at ∼13kDa could be detected, corresponding to incomplete cleavage at the C-terminal end. In contrast, even with prolonged incubation, SapA appeared as a series of fragments ranging from 9-21kDa in size (Fig. 2).

**Figure 2.**
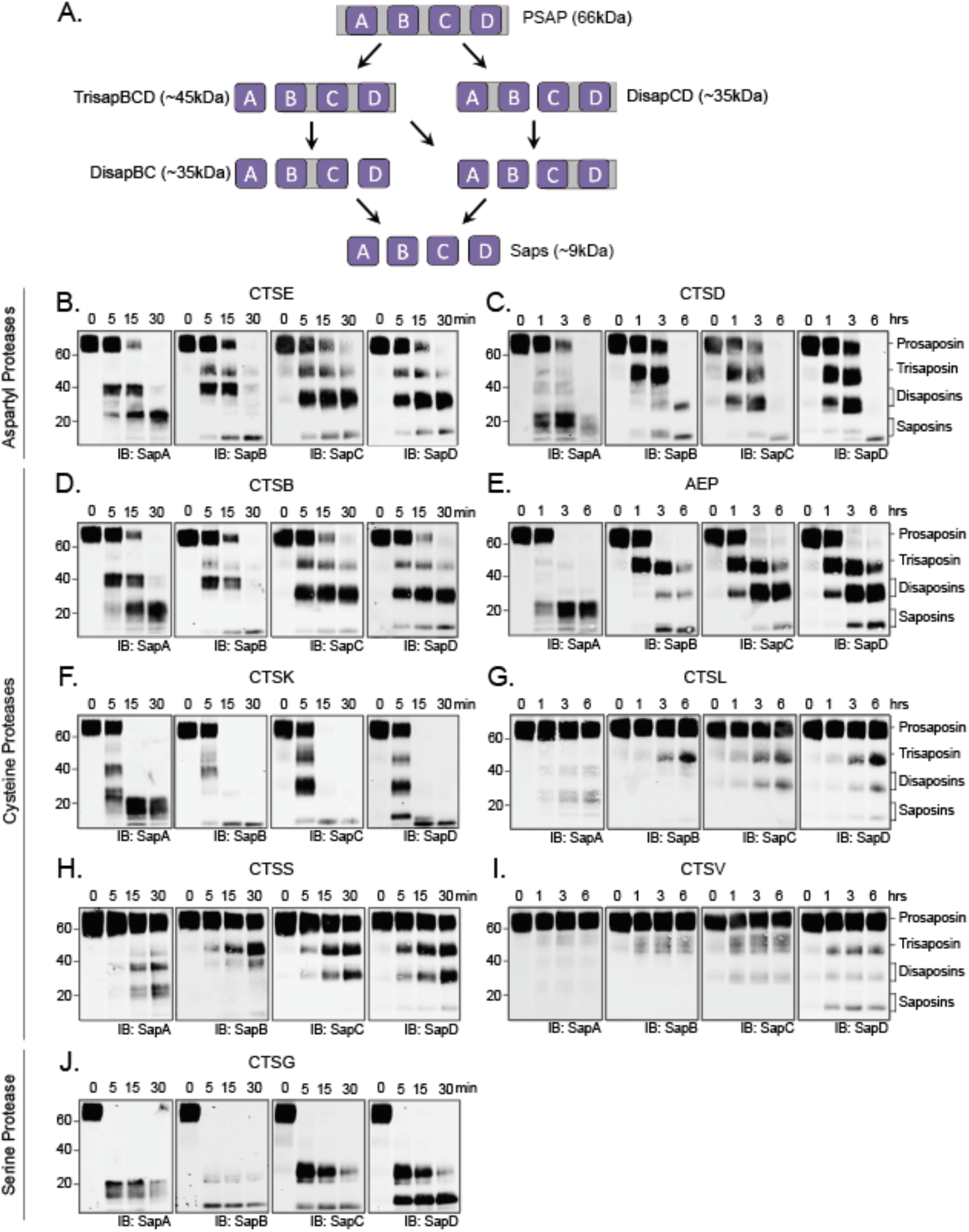
Saposins A-D are released by several cathepsins. *A*, Model of PSAP cleavage into saposins. Human recombinant prosaposin (PSAP, 400nM) was incubated at 37°C for either a 30 minute or 6 hour time course with various human recombinant cathepsins (400nM) at their optimal pH. *B-C*, Aspartyl proteases CTSD and CTSE, *D-I*, cysteine proteases CTSB, AEP/LGMN, CTSK, CTSL, CTSS, and CTSV, and *J*, serine protease CTSG were tested. Samples were run on western blot and probed with α-SapA, α-SapB, α-SapC, α-SapD antibodies. All enzymes were tested at pH of 4.5, with the exception of CTSG, which was tested at its optimal pH of 7.4.

In the aspartyl family of proteases, both cathepsins E and D produce all four saposins (Fig. 2B-C). CTSE completely digested PSAP by 30 minutes, while CTSD took 6 hours. In the cysteine family of proteases, CTSB, CTSK, CTSL, CTSS and AEP/LGMN are able to produce all four saposins (Fig. 2D-H), albeit with different relative efficiencies. On one end, CTSK was able to fully cleave PSAP into individual saposins within 15 minutes, whereas CTSL produced only minimally detectable amounts of individual saposins after 6 hours. Uniquely, CTSV mostly produced a singular individual saposin, SapD, and fragment abundance did not increase after the 1-hour time-point, indicating that prolonging incubation past 6 hours would be unlikely to change the results (Fig. 2I). Lastly, and most strikingly, the serine family protease CTSG was the most successful cathepsin tested. It was able to completely digest PSAP within 5-minutes, and produced all four saposins (Fig. 2J).

These experiments demonstrated that most cathepsins that cleave PSAP were able to produce all four saposins *in vitro*, though with variable efficacy. We also found that for all proteases tested, SapA was the most abundant or only saposin present at the shortest incubation length (Fig. 2), consistent with the previous observation that SapA is the first saposin released from PSAP [43]. Accordingly, the remainder of the protein, TrisaposinBCD (45kDa), was also present at high quantities at these time points. The first cleavage of PSAP also occasionally occured between SapB and SapC, generating DisaposinAB (35kDa) and DisaposinCD (35kDa). SapB and SapC were also found at the shortest tested incubation lengths for some cathepsins (Fig. 2). In contrast, none of the cathepsins tested cleaved full-length PSAP between SapC and SapD, as indicated by the complete lack of TrisaposinABC and the relative delay of the presence of SapD (17kDa), which must be digested from the downstream products TrisaposinBCD and DisaposinCD (Fig. 2). DisaposinCD abundance persistsed longer than DisaposinAB abundance, and SapC (17kDa) often is the slowest saposin to be produced.

To further understand the diversity of PSAP cleavage products produced by each cathepsin, we used the PROSPER database [62] to predict intersaposin cleavage sites *in silico* (Fig. 3, Supp. Table 1). Of the cathepsins we found to cleave PSAP, all were available in the database except CTSV and AEP. Similar to our results *in vitro*, cathepsins D, E, B, K and L were predicted to cleave in all intersaposin regions. While our data indicate cathepsins S and G also cleave all intersaposin regions, the PROSPER analysis only predicted CTSS to cleave between SapA-SapB and SapC-SapD, and CTSG to cleave in the intersaposin region between SapA-SapB and SapB-SapC.

**Figure 3.**
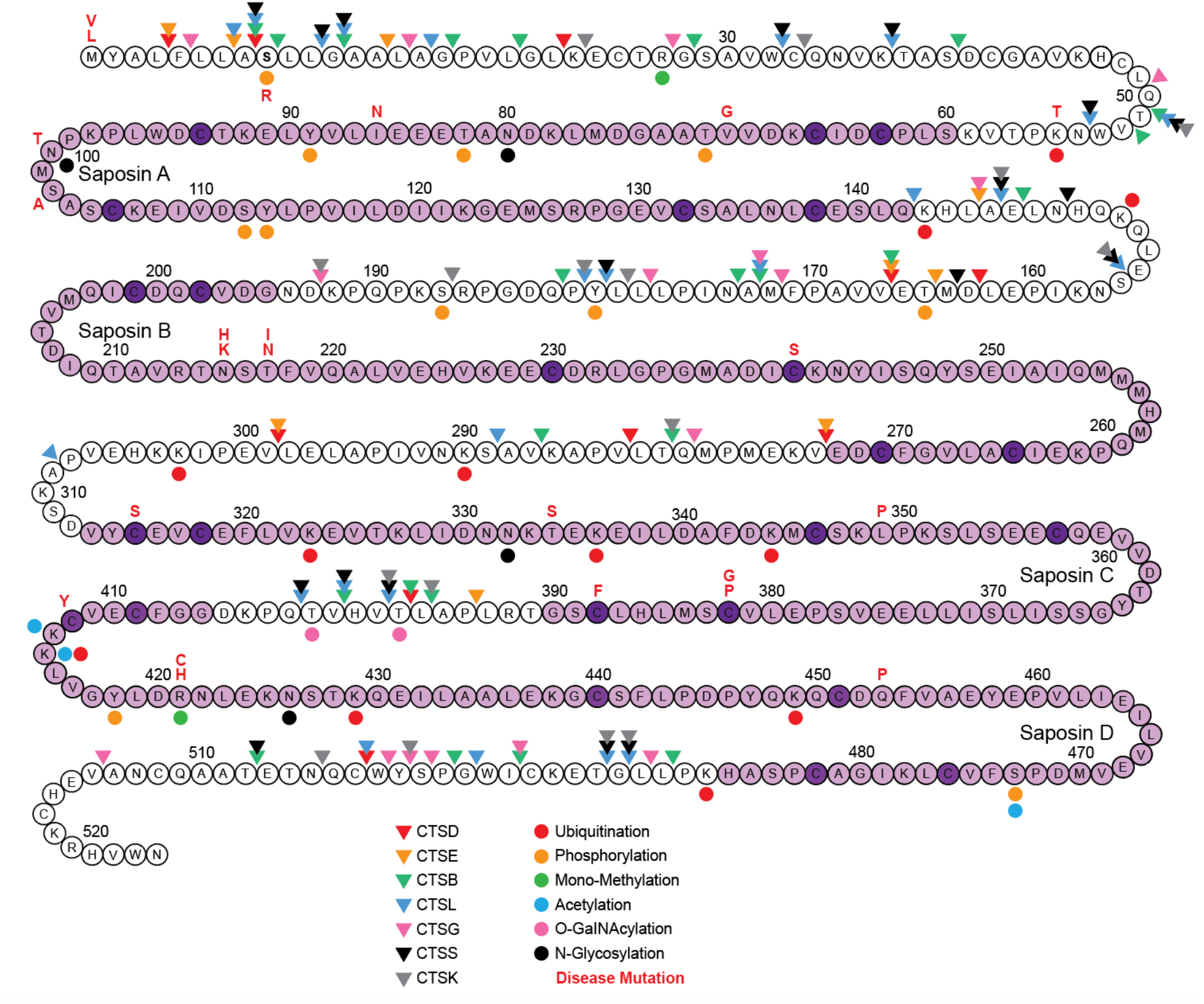
Map of PSAP predicted cleavage sites, post-translational modifications, and disease-causing mutations. A string-of-pearls model of PSAP. Cathepsin cleavage sites were predicted *in silico* using the PROSPER database [62]. For a list of predicted cleavage sites, see Supplemental Table 1. Post-translational modifications (PTMs) and disease-causing mutations were identified via literature review. For a list of predicted PTM and mutation sites, see Supplemental Table 2.

Together, these data indicate that some saposins are more readily produced than others. Considering the unique role for each saposin in sphingolipid metabolism, the ability of various cathepsins to produce less abundant saposins, like SapC, may be critical to the maintenance of sphingolipid homeostasis.

### Progranulin regulates prosaposin cleavage via Cathepsin D

Considering the importance of saposins in the regulation of sphingolipidase activity, we sought to identify upstream regulators of PSAP cleavage by cathepsins. The PSAP binding partner progranulin (PGRN) has been shown to regulate saposin abundance [46, 63] and it has been recently implicated in disease-related dysfunction in lipid homeostasis [49, 50]. PGRN has been shown to directly bind to and regulate the subcellular localization of PSAP [45, 46]. Further, PGRN and multi-granulin fragments (MGFs) have been shown to directly bind to and regulate the activity of CTSD [52-54]. Based on this, we hypothesized that PGRN may play a role in PSAP cleavage via modulation of CTSD activity.

As another lysosomal pro-protein, PGRN is similarly cleaved into MGFs and individual granulins (Fig 4A) by a number of cathepsins [51]. We tested three MGFs, pG, BAC, and CDE for their ability to regulate PSAP cleavage by CTSD. To test the effect of PGRN and MGFs on CTSD-mediated cleavage of PSAP, we incubated PSAP with CTSD for 6 hours with PGRN or MGFs and measured saposin production. Interestingly, incubation of CTSD with PGRN decreased PSAP cleavage in vitro (Fig. 4B-E). Further, while incubation of CTSD with pG did not change PSAP cleavage, incubation with BAC or CDE increased the rate of PSAP cleavage, with CDE having the largest effect (Fig. 4B-E). This suggests a novel mechanism for PGRN in the regulation of PSAP, positioning it as an upstream regulator of PSAP cleavage and saposin production.

**Figure 4.**
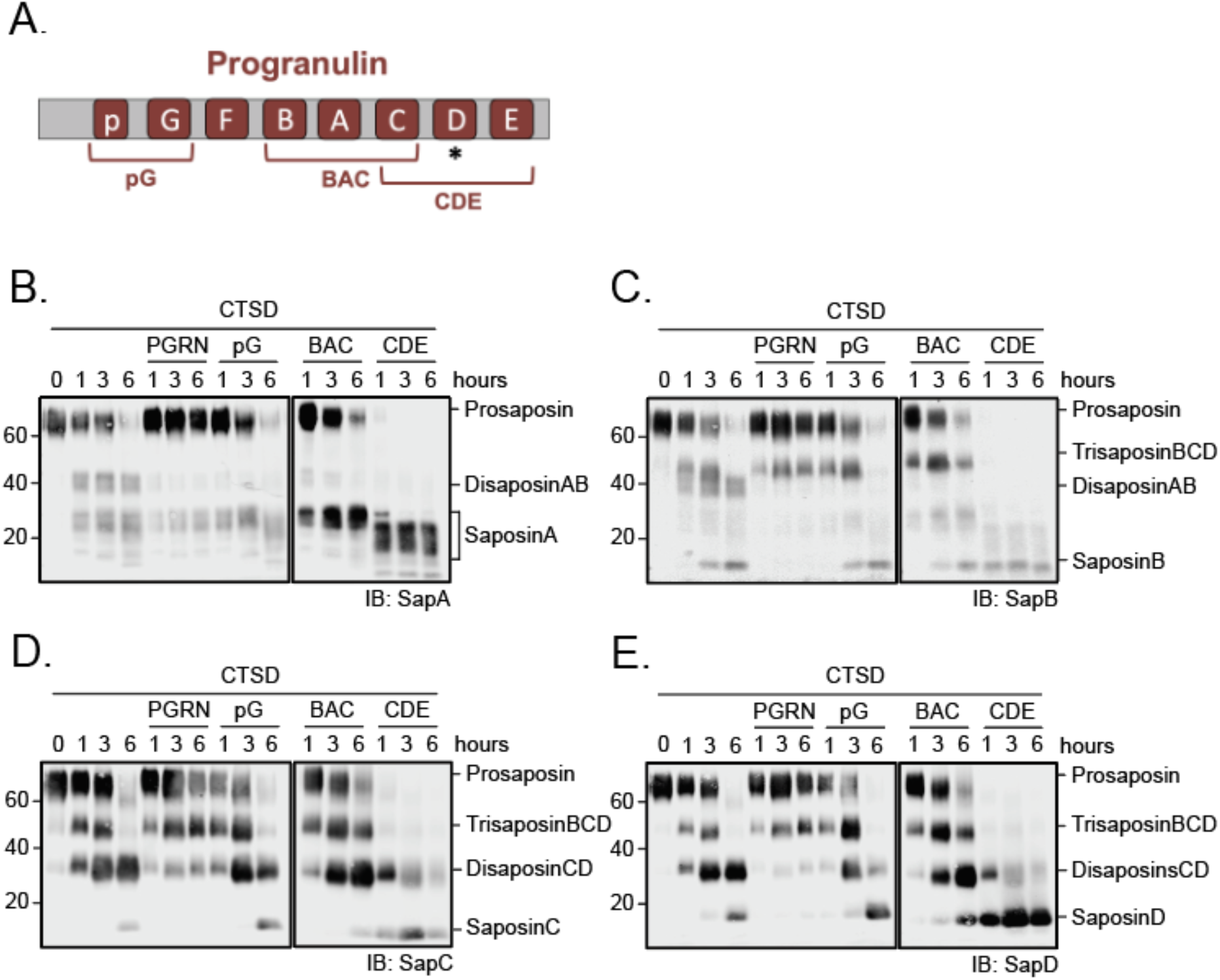
Progranulin and multi-granulin fragments influence the rate of PSAP cleavage and production of saposins A-D. *A*, Model of progranulin (PGRN) showing full length, and the three multi-granulin fragments (MGFs) pG, BAC, and CDE. * indicates the PSAP binding site on PGRN. *B-E*, Human recombinant prosaposin (PSAP,) was incubated at 37°C for a 6 hour time course with CTSD (400nM) and full-length PGRN, pG, BAC, or CDE (400nM) at pH 4.5. Samples were run on western blot and probed with α-SapA, α-SapB, α-SapC, α-SapD antibodies.

## DISCUSSION

In this study, we surveyed lysosomal cathepsins to determine which proteases are capable of PSAP cleavage, the pH-dependence of their activity, and their ability to produce specific saposin species. Considering the key role of saposins in sphingolipid metabolism, lysosomal storage disorders and age-related neurodegenerative diseases, better understanding of the proteases responsible for saposin production has potential therapeutic implications. Previously, the only known proteases to cleave PSAP into saposins were cathepsins D [39] and B [40]. Here, we showed that multiple cathepsins (E, K, L, S, V, G and AEP/LGMN) can process PSAP into saposins as well. Moreover, we found that nearly all of the proteases which process PSAP do so predominantly at the lysosomal pH of 3.5-4.5, suggesting that the majority of PSAP processing occurs in the lysosome, consistent with the canonical role of saposins in modulating lysosomal shingolipidase activity. Surprisingly, our data indicate that PSAP cleavage can also occur at a neutral pH 7.4 by cathepsins G, B, K, and S. All four of these cathepsins can be secreted [56][59-62], suggesting that they may also cleave PSAP extracellularly. Prosaposin is secreted into extracellular space both within the CNS and also by several tissues in the periphery. While extra-cellular prosaposin exerts effects on G-protein coupled receptors signaling and ERK phosphorylation[64, 65], extracellular saposins are relatively understudied.

Our data suggests that many cathepsins can uniquely contribute to the generation of saposins A-D, and thus may influence sphingolipid metabolism and relative saposin levels. Accordingly, each saposin-linked lysosomal storage disorder is associated with an overlapping but non identical set of cathepsins [e.g, Neimann-Pick (Saposin A, Cathepsins D, B, L and S), Metachromatic Leukdystrophy (Saposin B, Cathepsins D and L), and Gaucher’s disease (Saposin C, Cathepsin D, B, K, and S)] [66-76]. Thus, relative levels of individual cathepsins may have nuanced but important effects on lysosomal PSAP cleavage and downstream development of disease.

Our lab and others have previously shown that CTSD activity is directly regulated by PGRN and multi-granulin fragments (MGFs) [52-54]. Here, we showed that PGRN and MGF regulation of CTSD activity directly affects the rate of saposin production. First, we found that full-length PGRN does not increase PSAP processing but rather seems to slow saposin production. While this finding was unexpected, it may be due to steric hindrance or structural alternations in the proteins. On the other hand, while the MGFs containing pG and BAC have small to negligible effects on the rate of saposin production, the MGF containing CDE increases the rate of saposin production considerably. The granulin D domain of PGRN has been reported to directly bind the intersaposin region beween SapB and SapC in PSAP [45, 46], suggesting that the direct binding of CDE to PSAP may be involved in the increased cleavage. We found that CTSD, despite being the most well-studied cathepsin to process PSAP, is among the least efficient. Thus, CDE’s considerable acceleration of its cleavage may have substantial effects *in vivo*. Our study is the first to report direct influence of PGRN and MGFs on the cleavage of PSAP, providing a biochemical mechanism that supports previous studies reporting that PGRN expression affects saposin abundance in cells [46, 63] and disrupting PGRN function leads to in disease-related dysfunction in lipid homeostasis [49, 50, 77]. Future studies that characterize the binding interaction of CTSD, PGRN/MGFs, and PSAP could provide further insight to the importance of this regulation *in vivo*.

While these results show that most cathepsins can independently generate all saposins *in vitro*, they do not directly represent how the cathepsins work in *in vivo*. Additionally, these studies lack potential PSAP binding partners and co-factors that can modulate protease activity *in vivo*, such as peptide inhibitors like cystatins and aspartins. However, this study is the first and most comprehensive of its kind to show the potential for many lysosomal proteases to process PSAP, implicating each of these cathepsins in the downstream regulation of sphingolipid metabolism and homeostasis.

In summary, our findings indicate that the processing of PSAP into saposins is subject to intricate regulatory mechanisms. These mechanisms involve protease-specific cleavage sites within the intersaposin regions and different cleavage rates based on pH environments. Notably, our study reveals a novel role for the PSAP binding partner PGRN in directly regulating saposin production. Future investigations may delve into the physiological significance of PSAP cleavage regulation by these proteases *in vivo*, particularly within the context of their cell-type specific expression. Considering the key role of PSAP and saposins in sphingolipid metabolism and the existence of genetic mutations in PSAP associated with neurodegenerative diseases in humans [2-7], it is clear that maintaining optimal levels of PSAP and saposins is crucial to maintain proper lysosomal function and sphingolipid homeostasis. With this study, we have identified PGRN, MGFs, and multiple cathepsins as key contributors to the production of saposins, which could present novel opportunities to modulate saposin levels in disease.

## EXPERIMENTAL PROCEDURES

### In vitro cleavage assays

400nM of recombinant human prosaposin (Abcam #167924) was incubated with or without 400nM of each protease and with or without recombinant human progranulin (Fisher #2420-PG) or recombinant human multi-granulin fragments pG (Mybiosources #2012256), BAC (Mybiosources #2011118), or CDE (Mybiosources #2018332). Reactions at pH 3.5 were performed in 100mM sodium citrate buffer, reactions at pH 4.5 and 5.5 were performed in 50mM sodium acetate, and reactions at pH 7.4 were performed in 100mM phosphate buffer saline (PBS). 20µL reactions were incubated with 1mM EDTA and 2mM DTT for 60 minutes in a 37° C water bath. Protease activity was stopped by adding 7.5 µl of NuPAGE 4X LDS (Fisher #NP0007), 3 µl of 10X reducing agent (Fisher #NP0009) and denatured for 10 minutes at 80°C. The samples were run on precast NOVEX 4-12% Bis-Tris gels (Fisher #NP0321PK2) using NOVEX-NuPAGE MOPS buffer (Fisher #NP0001). The gel was then either fixed in 40% ethanol and 10% acetic acid for silver staining or transferred onto nitrocellulose membranes for western blotting analysis.

### Recombinant proteases

Cathepsin A (R&D # 1049-SE), Cathepsin B (Millipore #219364), Cathepsin C (R&D #1071-CY), Cathepsin D (R&D #1014-AS), Cathepsin E (R&D #1294-AS), Cathepsin F (Abcam #157039), Cathepsin G (Millipore #219873), Cathepsin H (R&D #7516-CY-010), Cathepsin K (Millipore #219461), Cathepsin L (Millipore #219402), Cathepsin O (Abcam # ab158237), Cathepsin S (R&D #1183-CY), Cathepsin V (R&D #1080-CY), Cathepsin W (Abcam # ab158238), Cathepsin X (R&D #934-CY), Asparagine Endopeptidase/Legumain (R&D #2199-CY).

### Activation of proteases

For Cathepsin A and Cathepsin C activation, 4μM of each protease was incubated with 200nM of Cathepsin L at room temperature for 60 minutes. After 60 minutes, 50μM of benzyloxycarbonyl FY(t-Bu)-DMK (Sigma #219427), an irreversible, highly specific inhibitor of Cathepsin L was added to quench cathepsin L activity. Once activated, Cathepsin A and C were used in the experiments outlined. Similarly, for Cathepsin H activation, 4μM of Cathepsin H was incubated with 500nM of thermolysin (R&D #3097-ZN) at room temperature for 3 hours. After 3 hours, 1 mM of Phosphoramidon (Tocris Bioscience #6333), a specific Thermolysin inhibitor, was added to quench Thermolysin activity. Once activated, cathepsin H was used in the experiments outlined. All other recombinant proteases are active, as shown by the vendors.

### Silver stains

Silver staining was performed according to manufacturer’s instructions with SilverQuest silver staining kit (Thermo #LC6070).

### Western blots

Western blots were performed on nitrocellulose membranes. Membranes were blocked with Odyssey buffer (Li-cor, #927-50010) for 1-2hrs and subsequently blotted with 1:1000 anti-prosaposin pAb (Abcam #180751), anti-saposin A pAb (Proteintech #18396-1-AP), anti-saposin B (Proteintech #18397-1-AP), anti-saposin C (Proteintech #18398-1-AP), and/or anti-saposin D (Proteintech #18423-1-AP) for 1hr. All prosaposin and saposin antibodies were raised in rabbit. Antibodies were validated by Abcam and Proteintech, respectively. Li-cor fluorescent secondary antibodies were used at 1:10000 dilution and incubated for 1hr. Western blots were imaged using an Odyssey CLx imager. As a positive control, I tested the antibodies against recombinant human prosaposin (Abcam #167924) alone and observed the expected band sizes.

