## Supplemental Tables for "Prosaposin is cleaved into saposins by multiple cathepsins in a progranulin-regulated fashion"

Supporting Information:

1. Supplemental Table 1 – Page S1
2. Supplemental Table 2 – Page S2

Supplemental Table 1.

| | CTSD | CTSE | CTSB | CTSL | CTSG | CTSS | CTSK |
|---|---|---|---|---|---|---|---|
| Before SapA 1-59 | 5: MYAL†FLLA<br>9: FLLA†SLLG<br>23: VLGL†KECT | 5: MYAL†FLLA<br>8: LFLL†ASLL | 5: LGAA†LAGP<br>9: FLLA†SLLG<br>10: LLAS†LLGA<br>13: SLLG†AALA<br>15: LGAA†LAGP<br>18: ALAG†PVLG<br>21: GPVL†GLKE<br>29: CTRG†SAVW<br>41: KTAS†DCGA<br>51: HCLQ†TVWN<br>52: CLQT†VWNK | 8: LFLL†ASLL<br>9: FLLA†SLLG<br>12: ASLL†GAAL<br>13: SLLG†AALA<br>17: AALA†GPVL<br>33: SAVW†CQNV<br>38: QNVK†TASD<br>51: HCLQ†TVWN<br>54: QTVW†NKPT | 6: YALF†LLAS<br>16: GAAL†AGPV<br>28: ECTR†GSAV<br>50: KHCL†QTVW | 9: FLLA†SLLG<br>12: ASLL†GAAL<br>13: SLLG†AALA<br>33: SAVW†CQNV<br>38: QNVK†TASD<br>51: HCLQ†TVWN<br>54: QTVW†NKPT | 24: LGLK†ECTR<br>34: AVWC†QNVK<br>51: HCLQ†TVWN |
| SapA 60-142 | 68: DICK†DVVT<br>88: EEIL†VYLE<br>90: ILVY†LEKT<br>91: LVYL†EKTC<br>112: EIVD†SYLP<br>119: PVIL†DIIK<br>120: VILD†IIKG<br>138: ALNL†CESL | 72: DVVT†AAGD<br>86: TEEE†ILVY<br>88: EEIL†VYLE<br>119: PVIL†DIIK<br>138: ALNL†CESL | 71: KDVV†TAAG<br>72: DVVT†AAGD<br>103: KPNM†SASC | 60: PTVK†SLPC<br>71: KDVV†TAAG<br>72: DVVT†AAGD<br>112: EIVD†SYLP<br>133: GEVC†SALN | 60: PTVK†SLPC<br>67: CDIC†KDVV<br>78: GDML†KDNA<br>88: EEIL†VYLE<br>114: VDSY†LPVI<br>136: CSAL†NLCE | 60: PTVK†SLPC<br>71: KDVV†TAAG<br>72: DVVT†AAGD<br>92: VYLE†KTCD<br>99: DWLP†KPNM<br>103: KPNM†SASC<br>104: PNMS†ASCK | 63: KSLP†CDIC<br>79: DMLK†DNAT<br>92: VYLE†KTCD<br>99: DWLP†KPNM<br>100: WLPK†PNMS<br>108: ASCK†EIVD<br>114: VDSY†LPVI<br>116: SYLP†VILD<br>117: YLPV†ILDI<br>139: LNLC†ESLQ |
| IntersapAB 143-194 | 163: IPEL†DMTE<br>167: DMTE†VVAP | 146: QKHL†AELN<br>165: ELDM†TEVV<br>167: DMTE†VVAP | 148: HLAE†LNHQ<br>167: DMTE†VVAP<br>173: APFM†ANIP<br>174: PFMA†NIPL<br>182: LLYP†QDGP | 143: ESLQ†KHLA<br>147: KHLA†ELNH<br>156: KQLE†SNKI<br>173: APFM†ANIP<br>180: PLLL†YPQD<br>181: LLLY†PQDG | 146: QKHL†AELN<br>172: VAPF†MANI<br>173: APFM†ANIP<br>178: NIPL†LLYP<br>193: PQPK†DNGD | 139: LNLC†ESLQ<br>147: KHLA†ELNH<br>150: AELN†HQKQ<br>156: KQLE†SNKI<br>164: PELD†MTEV<br>180: PLLL†YPQD | 147: KHLA†ELNH<br>156: KQLE†SNKI<br>179: IPLL†LYPQ<br>181: LLLY†PQDG<br>187: DGPR†SKPQ<br>193: PQPK†DNGD |
| SapB 195-273 | 202: CQDC†IQMV<br>208: MVTD†IQTA<br>219: NSTF†VQAL<br>223: VQAL†VEHV<br>226: LVEH†VKEE<br>253: SEIA†IQMM<br>257: IQMM†MHMQ<br>267: EICA†LVGF<br>268: ICAL†VGFC<br>271: LVGF†CDEV | 202: CQDC†IQMV<br>204: DCIQ†MVTD<br>205: CIQM†VTDI<br>219: NSTF†VQAL<br>221: TFVQ†ALVE<br>223: VQAL†VEHV<br>251: QYSE†IAIQ<br>253: SEIA†IQMM<br>257: IQMM†MHMQ<br>258: QMMM†HMQP<br>268: ICAL†VGFC | 207: QMVT†DIQT<br>218: TNST†FVQA<br>221: TFVQ†ALVE<br>222: FVQA†LVEH<br>258: QMMM†HMQP<br>267: EICA†LVGF<br>268: ICAL†VGFC<br>270: ALVG†FCDE | 214: TAVR†TNST<br>225: ALVE†HVKE<br>246: KNYI†SQYS<br>258: QMMM†HMQP<br>269: CALV†GFCD<br>270: ALVG†FCDE | 195: PKDN†GDVC<br>202: CQDC†IQMV<br>205: CIQM†VTDI<br>216: VRTN†STFV<br>223: VQAL†VEHV<br>249: ISQY†SEIA<br>255: IAIQ†MMMH<br>256: AIQM†MMHM<br>257: IQMM†MHMQ<br>258: QMMM†HMQP<br>259: MMMH†MQPK<br>268: ICAL†VGFC | 207: QMVT†DIQT<br>221: TFVQ†ALVE<br>225: ALVE†HVKE<br>246: KNYI†SQYS<br>258: QMMM†HMQP<br>259: MMMH†MQPK<br>269: CALV†GFCD<br>270: ALVG†FCDE | 199: GDVC†QDCI<br>204: DCIQ†MVTD<br>205: CIQM†VTDI<br>224: QALV†EHVK<br>237: LGPG†MADI<br>245: CKNY†ISQY<br>255: IAIQ†MMMH<br>257: IQMM†MHMQ<br>259: MMMH†MQPK<br>269: CALV†GFCD |
| IntersapBC 274-312 | 274: FCDE†VKEM<br>283: MQTL†VPAK<br>299: ALEL†VEPI | 274: FCDE†VKEM<br>299: ALEL†VEPI | 281: MPMQ†TLVP<br>282: PMQT†LVPA<br>287: VPAK†VASK | 289: AKVA†SKNV<br>309: HEVP†AKSD | 280: EMPM†QTLV | | 281: MPMQ†TLVP |
| SapC 313-390 | 320: EVCE†FLVK<br>321: VCEF†LVKE<br>322: CEFL†VKEV<br>329: VTKL†IDNN<br>434: QEIL†AALE<br>340: KEIL†DAFD<br>342: ILDA†FDKM<br>343: LDAF†DKMC<br>373: LSIL†LEEV<br>381: SPEL†VCSM | 317: VYCE†VCEF<br>321: VCEF†LVKE<br>322: CEFL†VKEV<br>329: VTKL†IDNN<br>343: LDAF†DKMC<br>354: PKSL†SEEC<br>360: ECQE†VVDT<br>370: SSIL†SILL<br>373: LSIL†LEEV<br>381: SPEL†VCSM<br>385: VCSM†LHLC<br>388: MLHL†CSGT | 363: EVVD†TYGS<br>383: ELVC†SMLH<br>387: SMLH†LCSG<br>388: MLHL†CSGT | 355: KSLS†EECQ<br>363: EVVD†TYGS<br>375: ILLE†EVSP<br>383: ELVC†SMLH<br>387: SMLH†LCSG<br>389: LHLC†SGTR | 315: SDVY†CEVC<br>321: VCEF†LVKE<br>322: CEFL†VKEV<br>329: VTKL†IDNN<br>354: PKSL†SEEC<br>365: VDTY†GSSI<br>373: LSIL†LEEV<br>386: CSML†HLCS | 355: KSLS†EECQ<br>362: QEVV†DTYG<br>363: EVVD†TYGS<br>375: ILLE†EVSP<br>383: ELVC†SMLH<br>387: SMLH†LCSG<br>389: LHLC†SGTR | 330: TKLI†DNNK<br>355: KSLS†EECQ<br>374: SILL†EEVS<br>375: ILLE†EVSP<br>387: SMLH†LCSG<br>389: LHLC†SGTR |
| IntersapCD 391-405 | 397: LPAL†TVHV | 394: GTRL†PALT | 396: RLPA†LTVH<br>397: LPAL†TVHV<br>400: LTVH†VTQP | 398: PALT†VHVT<br>400: LTVH†VTQP<br>402: VHVT†QPKD | | 398: PALT†VHVT<br>400: LTVH†VTQP<br>402: VHVT†QPKD | 396: RLPA†LTVH<br>398: PALT†VHVT |
| SapD 406-486 | 409: DGGF†CEVC<br>447: LPDP†YQKQ<br>455: CDQF†VAEY<br>458: FVAE†YEPV<br>463: EPVL†IEIL<br>466: LIEI†LVEV<br>467: IEIL†VEVM<br>469: ILVE†VMDP<br>475: DPSF†VCLK<br>477: SFVC†LKIG<br>478: FVCL†KIGA | 411: GFCE†VCKK<br>424: DRNL†EKNS<br>432: TKQE†ILAA<br>434: QEIL†AALE<br>443: GCSF†LPDP<br>455: CDQF†VAEY<br>463: EPVL†IEIL<br>465: VLIE†ILVE<br>467: IEIL†VEVM<br>469: ILVE†VMDP<br>475: DPSF†VCLK<br>477: SFVC†LKIG<br>478: FVCL†KIGA | 415: VCKK†LVGY<br>418: KLVG†YLDR<br>436: ILAA†LEKG<br>437: LAAL†EKGC<br>442: KGCS†FLPD<br>454: QCDQ†FVAE<br>457: QFVA†EYEP<br>469: ILVE†VMDP<br>478: FVCL†KIGA | 417: KKLV†GYLD<br>425: RNLE†KNST<br>445: SFLP†DPYQ<br>449: DPYQ†KQCD<br>457: QFVA†EYEP<br>469: ILVE†VMDP<br>480: CLKI†GACP<br>485: ACPS†AHKP | 411: DGGF†CEVC<br>434: QEIL†AALE<br>444: CSFL†PDPY<br>454: QCDQ†FVAE<br>456: DQFV†AEYE<br>467: IEIL†VEVM<br>448: PDPY†QKQC<br>485: ACPS†AHKP | 418: KLVG†YLDR<br>425: RNLE†KNST<br>445: SFLP†DPYQ<br>457: QFVA†EYEP<br>471: VEVM†DPSF | 425: RNLE†KNST<br>442: KGCS†FLPD<br>449: DPYQ†KQCD<br>468: EILV†EVMD<br>479: VCLK†IGAC |
| After SapD 487-524 | 503: PSYW†CQNT | | 489: AHKP†LLGT<br>496: TEKC†IWGP<br>499: CIWG†PSYW<br>508: QNTE†TAAQ | 491: KPLL†GTEK<br>492: PLLG†TEKC<br>498: KCIW†GPSY<br>503: PSYW†CQNT | 490: HKPL†LGTE<br>496: TEKC†IWGP<br>501: WGPS†YWCQ<br>502: GPSY†WCQN<br>511: IWGP†SYWC<br>515: QCNA†VEHC | 491: KPLL†GTEK<br>492: PLLG†TEKC<br>508: QNTE†TAAQ | 491: KPLL†GTEK<br>492: PLLG†TEKC<br>501: WGPS†YWCQ<br>505: YWCQ†NTET |

**Supplemental Table 1.** ***In silico*** **predicted cleavage sites of PSAP.**

Supplemental Table 2.

| | Ubiquitination | Phosphorylation | Mono-Methylation | Acetylation | O-GalNAcylation | N-Glycosylation | Disease Mutations |
|---|---|---|---|---|---|---|---|
| Before SapA 1-59 | | S9 | R27 | | | | M1V M1L K55T |
| SapA 60-142 | | T72 S112 T82 Y112 Y88 | | | | N80 N101 | I86N N101T S103A |
| IntersapAB 143-194 | K143 K150 | T165 Y180 S187 | | | | | |
| SapB 195-273 | | | | | | | N215H T217N N215K C241S T217I |
| IntersapBC 274-312 | K290 K303 | | | | | | |
| SapC 313-390 | K323 K336 K344 | | | | | N426 | C315S C382G T334S C382P L249P C388F |
| IntersapCD 391-405 | | | | | T397 T401 | | |
| SapD 406-486 | K414 K429 K449 | Y418 S473 | R421 | K413 K414 S473 | | | C412Y Q452P R421C R421H |
| After SapD 487-524 | K487 | | | | | | |

**Supplemental Table 2. Post-translational modification sites and disease-causing mutations in PSAP.**
